# A regulatory role for the unstructured C-terminal domain of the CtBP transcriptional corepressor

**DOI:** 10.1101/2023.05.19.541472

**Authors:** Ana-Maria Raicu, Megha Suresh, David N. Arnosti

**Affiliations:** Cell and Molecular Biology Program, Michigan State University, East Lansing, MI; Department of Biochemistry and Molecular Biology, Michigan State University, East Lansing, MI

## Abstract

The C-terminal Binding Protein (CtBP) is a transcriptional corepressor that plays critical roles in development, tumorigenesis, and cell fate. CtBP proteins are structurally similar to alpha hydroxyacid dehydrogenases and feature a prominent intrinsically disordered region in the C-terminus. In the mammalian system, CtBP proteins lacking the C-terminal Domain (CTD) are able to function as transcriptional regulators and oligomerize, putting into question the significance of this unstructured domain for gene regulation. Yet, the presence of an unstructured CTD of ∼100 residues, including some short motifs, is conserved across Bilateria, indicating the importance of maintaining this domain over evolutionary time. To uncover the significance of the CtBP CTD, we functionally tested naturally occurring Drosophila isoforms of CtBP that possess or lack the CTD, namely CtBP(L) and CtBP(S). We used the CRISPRi system to recruit dCas9-CtBP(L) and dCas9-CtBP(S) to endogenous promoters to directly compare their transcriptional impacts *in vivo*. Interestingly, CtBP(S) was able to significantly repress transcription of the *Mpp6* promoter, while CtBP(L) was much weaker, suggesting that the long CTD may modulate CtBP’s repression activity. In contrast, in cell culture, the isoforms behaved similarly on a transfected *Mpp6* reporter gene. The context-specific differences in activity of these two developmentally-regulated isoforms suggests that the CTD may help provide a spectrum of repression activity suitable for developmental programs.

## INTRODUCTION

Eukaryotic transcription factors and cofactors are rich in unstructured domains; these proteins have a higher percentage of predicted intrinsically disordered regions (IDR) than the average protein [1]. Some of these IDRs have been shown to participate in specific transcriptional processes, sometimes through promoting the formation of phase separated condensates [2]. For example, the C-terminal domain (CTD) of RNA polymerase II, a well-studied IDR, is a platform for association of factors involved in capping, splicing, and polyadenylation [3]. The N-terminal IDR of the androgen receptor (AR) has also recently been found to be necessary for condensate formation and transcriptional activity on enhancers [4]. IDRs in a plethora of other transcriptional regulators are thought to similarly play roles related to gene regulation, although most remain understudied [2]. Recent high-throughput analyses to determine the function of IDRs across the eukaryotic proteome have uncovered important motifs, interacting partners, and putative gene regulatory functions of some of these uncharacterized IDRs [2]. However, the specific roles of many IDRs present in these factors are still unknown. Tools to probe the function of certain IDRs and examine their gene regulatory roles in a physiologically relevant context and in a developing organism are necessary to delineate mechanisms of gene regulation by these IDRs and start to assign *in vivo* functions to them.

A prominent IDR that has not been well-studied is the C-terminal Domain (CTD) of the C-terminal binding protein (CtBP). CtBP is a highly conserved transcriptional corepressor that plays a role in cell differentiation and apoptosis, and has been implicated in a variety of human cancers [5]. This IDR of approximately 100 amino acids is not necessary for oligomerization of CtBP and may not be necessary for gene regulation, putting into question the significance of the CtBP CTD [6, 7]. Yet, our recent study shows that the CtBP CTD is highly conserved across Bilateria, and despite possessing overall lower sequence conservation than other parts of the protein, it features conserved short linear motifs within this predicted unstructured domain [8]. A few lineages such as roundworms and flatworms have novel, derived CTD sequences that are predicted to form intrinsic structures of unknown function. However, the deep conservation in primary sequence, length, and unstructured property of the CtBP CTD in bilaterians suggests that this IDR plays an important role, perhaps in gene regulation [8].

Mammalian genomes encode the CtBP1 and CtBP2 paralogs, which play overlapping and non-redundant roles in regulating expression of genes involved in apoptosis, the epithelial to mesenchymal transition, and cell differentiation [9-12]. The CtBP1 and CtBP2 CTDs exhibit 50% sequence conservation, which is much lower than that of the central core dehydrogenase domain, which is used for oligomerization, NADH binding and *in vitro* dehydrogenase activity [6, 8]. Interestingly, CtBP isoforms without the CTD exist in certain tetrapods such as birds and amphibians [8]. Likewise, in Drosophila, the single CtBP gene encodes multiple splice forms, including short isoforms that lack the CTD (CtBP-short, or CtBP(S)) and others that retain the long CTD (CtBP-long, or CtBP(L)) [8, 13]. These two major isoforms differing in the retention or loss of the IDR are co-expressed in fly development [13]. Thus, Drosophila is an appropriate model system to test a possible role of the CtBP CTD in gene regulation and assign a function to this elusive IDR.

Previous work using GAL4-CtBP fusions in the Drosophila embryo demonstrated that the two isoforms have similar repressive effects on an *even-skipped*-lacZ reporter, and both isoforms individually rescue a *CtBP* null fly, albeit with some different phenotypes in the wing [14, 15]. Thus, the CtBP CTD does not seem to play an essential role for completion of developmental programs under laboratory conditions. The expression pattern of the two isoforms exhibit developmentally distinct profiles; CtBP(S) is expressed throughout development, while CtBP(L) is highly expressed in the embryonic stage [13]. The fact that short isoforms have been independently derived in other insects, such as Hymenoptera, and in other lineages in Bilateria suggests that expression of both isoforms is somehow important [8]. The strict evolutionary conservation in these lineages of CtBP isoforms with and without the CTD indicates that both are functional, but a role in gene expression has remained unclear, compelling us to directly compare the activity of CtBP isoforms *in vivo*.

Here, we have made use of precise genetic tools in Drosophila to probe the function of the fly CtBP isoforms, CtBP(L) and CtBP(S), to uncover the role of the C-terminal IDR in regulating gene expression. Specifically, we used the CRISPRi system in the developing fly to assess the function of chimeric dCas9-CtBP proteins targeted to diverse gene promoters *in vivo*. This method allowed us to compare the activity of the long and short isoforms on the same loci in fly wing tissue, and compare the results to those targeted to transfected reporters in cell culture. We found that when assessed on endogenous targets, CtBP(S) is a more potent repressor of the *Mpp6* promoter than CtBP(L), but that this difference in repression ability is not observed on a transiently transfected *Mpp6*-luciferase reporter. Thus, in some contexts, the disordered CTD seems to provide a regulatory function, but the difference observed between endogenous gene regulation and transient transfections raises the possibility that the effect may be chromatin-dependent. Additionally, gene promoters targeted here had differential sensitivity to CtBP recruitment, indicating a further level of regulatory specificity, in accord with recent high-throughput assays [16].

## RESULTS

### Creation of dCas9-CtBP chimeras to regulate gene expression

To investigate the function of the CtBP C-terminal IDR and differences in gene regulation by the CtBP(L) and CtBP(S) isoforms in Drosophila, we employed CRISPRi [17]. These two isoforms are created through alternative splicing, with CtBP(L) having a ∼130 amino acid domain and CtBP(S) a ∼30 amino acid domain, which only share the first 20 residues with one another [8]. We fused the coding sequence of each CtBP isoform to a nuclease dead Cas9 (dCas9) enzyme to recruit CtBP corepressors to target promoters using gene-specific guide RNAs (gRNA; **Figure 1A**). dCas9-CtBP(L) and dCas9-CtBP(S) are expressed in S2 cells, according to western blot (**Figure S1**).

**Figure 1.**
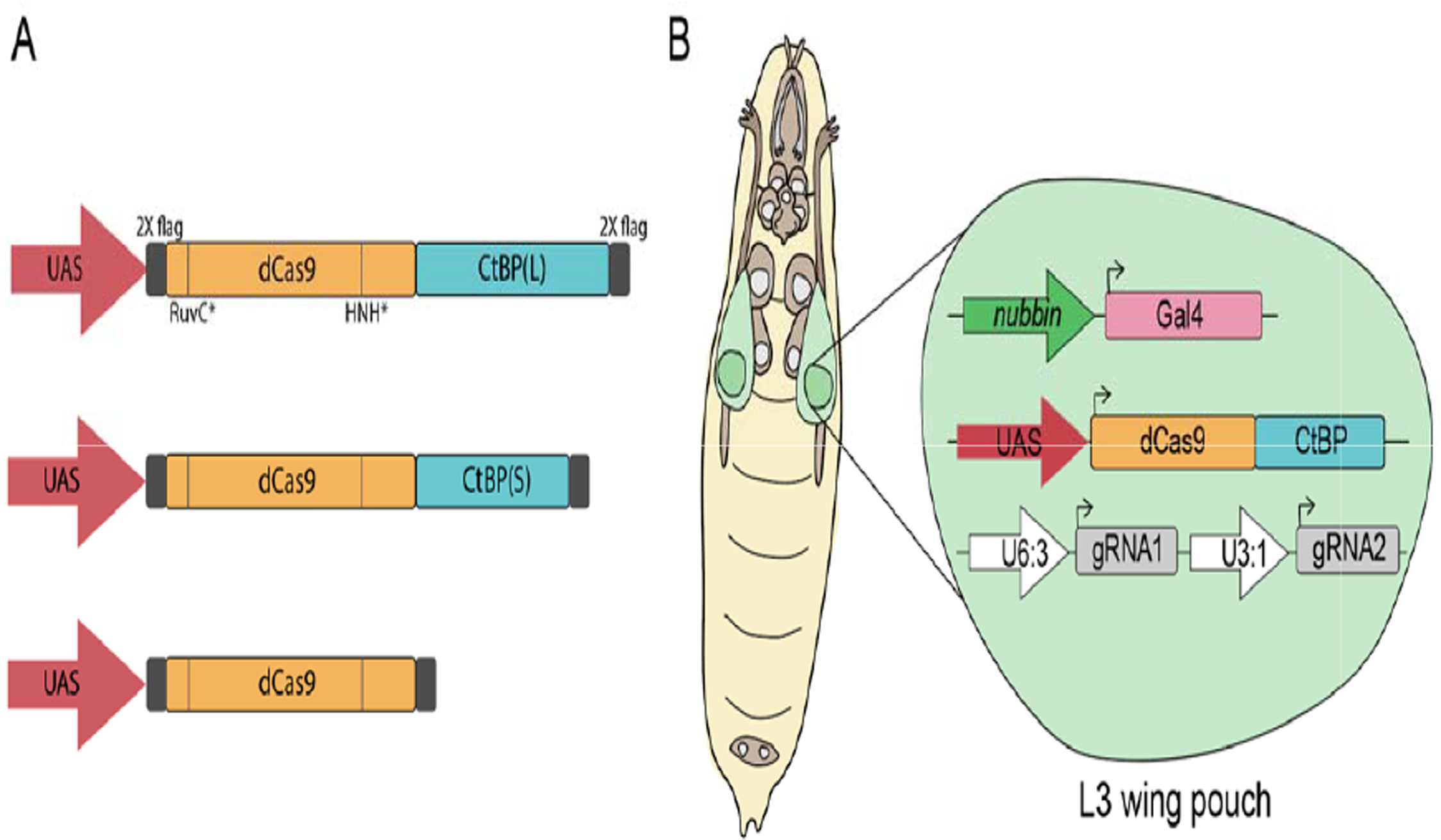
An *in vivo* system for targeting CtBP isoforms to gene promoters using CRISPRi. **A)** The fly CtBP(L) and CtBP(S) FLAG-tagged coding sequences were fused to the C-terminus of the *S. pyogenes* nuclease dead Cas9 (dCas9; D10A mutation in RuvC catalytic domain and H840A mutation in HNH catalytic domain), and placed under UAS expression. FLAG-tagged dCas9 was used as a negative control. Vertical lines in dCas9 represent the inactivating mutations. **B)** *Drosophila melanogaster* expressing three transgenes were generated for tissue-specific expression of dCas9-CtBP effectors using GAL4-UAS. Flies express dCas9-CtBP chimeras in the *nubbin* expression pattern (wing pouch of L3 wing discs), with ubiquitous expression of two tandem gRNAs designed to target a single gene’s promoter. Flies used in experiments express one copy of each of the three transgenes. gRNA flies were designed by the Drosophila Research and Screening Center and obtained from BDSC.

We expressed the chimeric proteins in the wing discs of third instar larvae (L3) using the *nubbin*-*GAL4* driver, which is predominantly expressed in the L3 wing pouch. Flies homozygous for both *nubbin-GAL4* and UAS:dCas9-CtBP were crossed to flies expressing two tandem gRNAs targeting diverse promoters (**Figure 1B)** [18]. These gRNAs were obtained as fly lines from the Bloomington Drosophila Stock Center (**Supplementary Table 1**). We previously tested dCas9-Rb chimeras in L3 discs, where we observed gene-specific effects after targeting ∼30 different gene promoters; here, we targeted many of the same promoters with the CtBP isoforms (**Supplementary Table 1**) [19].

The epithelial cells of the developing wing are a highly sensitive tissue that has been used to measure developmental perturbation of a number of regulatory pathways. To screen for genetic effects after targeting chimeras in cells of the L3 wing discs, we allowed the flies expressing the three transgenes to grow to adulthood, and then assessed adult wing phenotypes from targeting each promoter, as has been previously done in Drosophila CRISPR activation screens [20]. We note that the *nubbin*-*GAL4*>UAS:dCas9-CtBP flies crossed to a non-targeting gRNA control fly line (QUAS) produced mild wing phenotypes, consisting chiefly of supernumerary bristles (**Figure 2A**). We presume that ectopic CtBP, even when fused to dCas9, may interact with diverse endogenous CtBP binding sites on the genome, leading to these mild phenotypes. The QUAS control gRNAs used here did not produce phenotypes with dCas9-Rb corepressor fusions tested previously, so the phenotypic effect is CtBP-specific [19].

**Figure 2.**
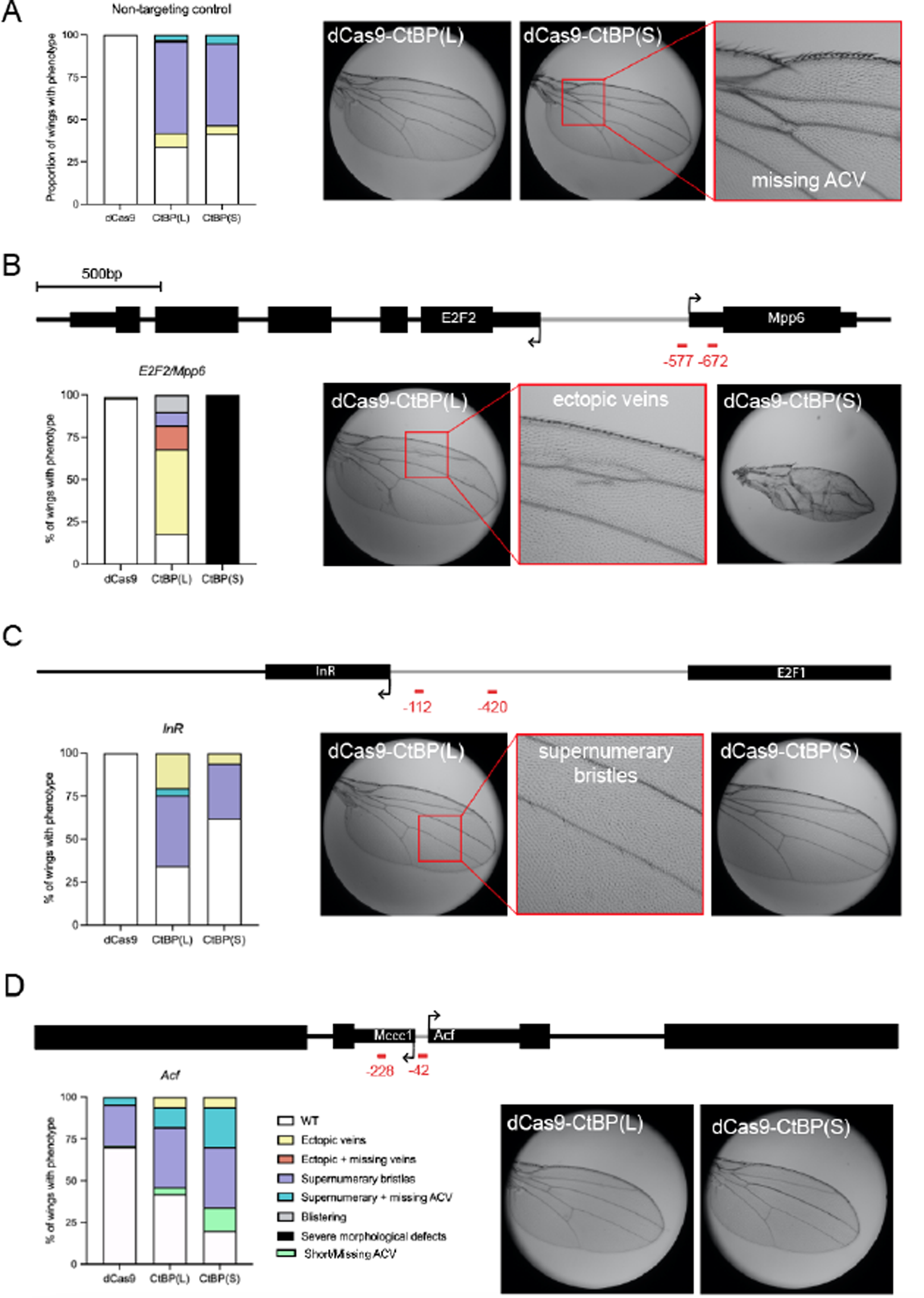
Targeting CtBP(S) and CtBP(L) to promoters leads to diverse phenotypic effects. Black arrows indicate the TSS, and red lines indicate gRNA binding sites relative to the target gene’s TSS. **A)** Using a non-targeting control gRNA (QUAS), expression of one copy of dCas9-CtBP effectors leads to >50% of adult wings with mild phenotypes such as supernumerary bristles. Legend is in panel D. ACV indicates Anterior Cross Vein. **B)** Targeting the *E2F2/Mpp6* bidirectional promoter leads to severe morphological defects observed only with CtBP(S) targeting, with milder effects caused by CtBP(L). gRNA positions are relative to the *E2F2* TSS, as designed by the Drosophila Research and Screening Center. **C)** Targeting the *InR* promoter leads to phenotypes similar to the QUAS non-targeting control, suggesting little or no specific effect on this promoter. **D)** Targeting the *Acf* promoter leads to mild phenotypes, some of which are also observed with dCas9 alone at lower frequency. CtBP isoforms lead to a higher penetrance of phenotypes than dCas9. For all crosses, ∼100 wings from ∼50 adults were used for analysis.

### Diverse effects of CtBP isoforms

We recruited CtBP(L) and CtBP(S) to a number of promoters, with specific effects observed only on a few promoters (**Supplementary Table 1**). Here, we detail the effects of targeting the *E2F2/Mpp6* bidirectional promoter, the insulin receptor (*InR*) promoter, and the promoter of *Acf*, a nucleosome remodeling subunit (**Figure 2**). Targeting CtBP(S) to the divergent *E2F2/Mpp6* promoter produced small wings with severe morphological defects, similar to that seen with dCas9-Rb fusions (**Figure 2B**) [19]. Intriguingly, CtBP(L) did not produce this phenotype, but instead produced much milder effects, including wings with ectopic veins and supernumerary bristles (**Figure 2B**). dCas9 alone did not produce any phenotypic effect, indicating that the observed phenotypes are CtBP-specific. The clear difference between targeting the long and short isoforms on this promoter suggests that the long CTD may inhibit CtBP’s gene regulatory activities.

The strong CtBP(S) effect is only seen when using two gRNAs; recruitment using the individual gRNAs at the same locus produced milder effects, such as ectopic veins seen with the CtBP(L) isoform when both gRNA were used (**Figure S2**). Interestingly, the number of wings with supernumerary bristles was less than that observed for the non-targeting control QUAS gRNA. We speculate that nonspecific CtBP overexpression effects are suppressed by targeting the chimeric protein to specific DNA locations using these single gRNAs.

Targeting the *InR* promoter produced adult wings with mild phenotypes, similar to those produced with the non-targeting QUAS gRNA control, so this effect is difficult to distinguish from a mild overexpression phenotype rather than specific *InR* targeting (**Figure 2C**). Clearly, positioning the CtBP chimeras near the *InR* transcriptional start site does not strongly affect the wing, although we know that positioning dCas9-Rb chimeras at this promoter does impact development and transcription [19]. This distinct effect is consistent with CtBP promoter selectivity, a property illustrated from recent high-throughput assays [16].

Recruitment to the *Acf* promoter region generated a different spectrum of phenotypes. In this case, a significant proportion of wings from the dCas9 control cross showed supernumerary bristles--evidence that dCas9 alone can disrupt gene function in certain locations. Notably, the position of one of the gRNAs used here is 3’ of the initiation site for the divergently transcribed *Mccc1* gene, a position from which transcriptional inhibition is possible by dCas9 [21]. Over and above the background of this dCas9 effect, the CtBP fusions had unique, specific effects, with CtBP(S) causing a larger proportion of wings to be affected (80%) than CtBP(L) (60%; **Figure 2D**). Results from these targeted promoters indicate that CtBP exhibits gene-specific effects, and that in some cases, CtBP(S) has a more pronounced effect than CtBP(L).

### CtBP(S) is a more potent transcriptional repressor of *Mpp6* than CtBP(L)

Given the noticeable difference in wing phenotype as a result of targeting the two CtBP isoforms to the *E2F2/Mpp6* shared promoter, we measured transcript levels of both of these genes in the L3 wing disc using RT-qPCR. The two gRNAs targeting *E2F2/Mpp6* bind at -577 and -672 relative to the *E2F2* TSS, and at -18 and +57 relative to the *Mpp6* TSS (**Figure 3A**). CtBP(S) targeting led to ∼50% repression of the *Mpp6* gene, whereas CtBP(L) effects were significantly weaker (∼25%) and statistically indistinguishable from those of dCas9 alone (**Figure 3C**). Effects on *E2F2* were more modest, with only ∼10% decrease in *E2F2* expression resulting from targeting by dCas9 and dCas9-CtBP(L), and no statistically significant change after targeting with dCas9-CtBP(S) (**Figure 3B**). The greater impact on *Mpp6* is likely a reflection of the inherent short-range of action of many transcriptional repressors and corepressors, which may influence chromatin structure over a span of a nucleosome [19, 22].

**Figure 3.**
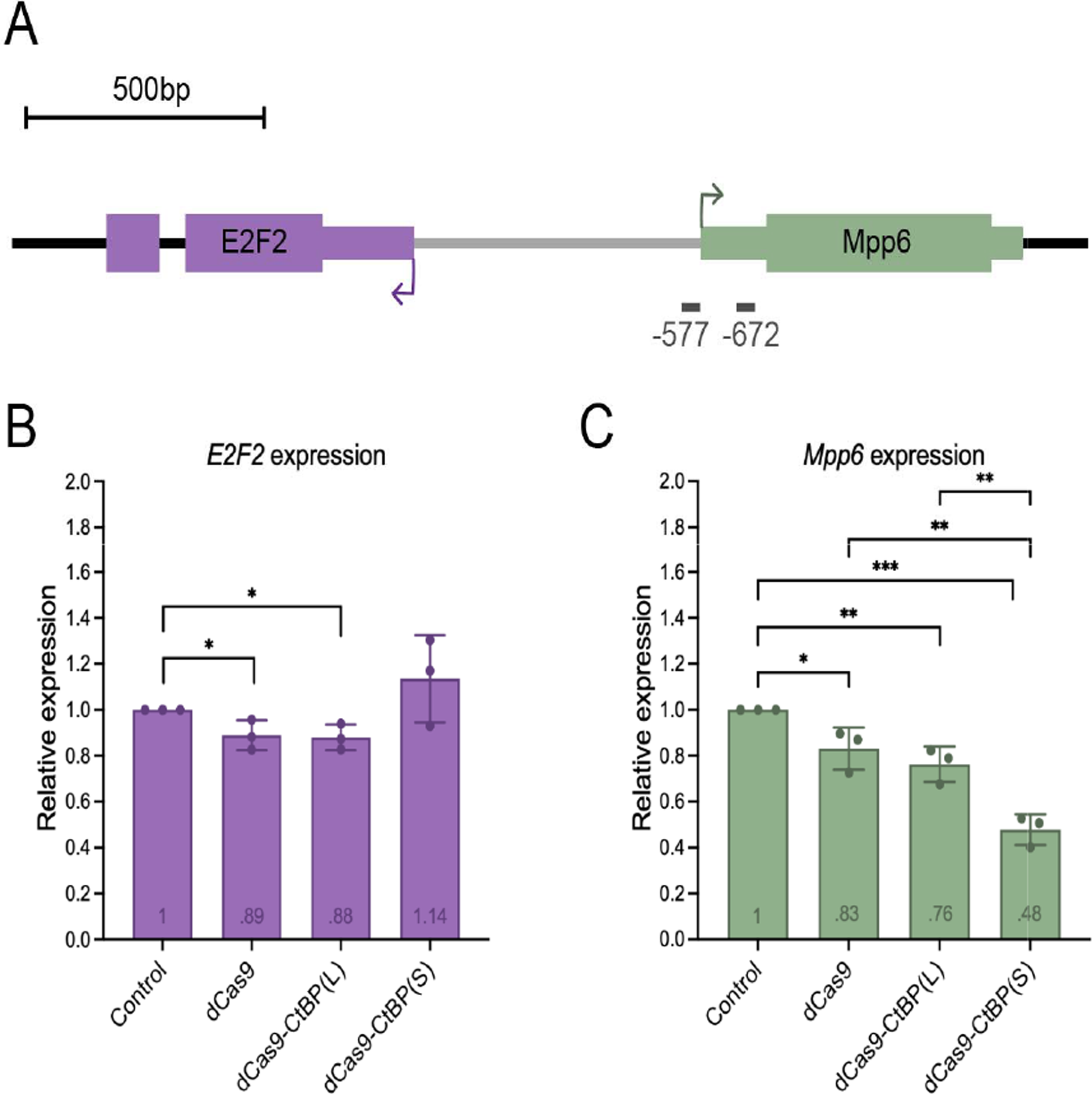
CtBP(S) is a more potent repressor of *Mpp6* than CtBP(L) in wing discs. **A)** Schematic of the *E2F2/Mpp6* bidirectional promoter, with the two tandem gRNAs indicated in gray (-577 and -672, positions relative to indicated *E2F2* TSS). **B)** Targeting dCas9 and dCas9-CtBP(L) led to modest repression (∼10%) of *E2F2*. Targeting CtBP(S) led to no statistically significant changes in *E2F2* expression. **C)** dCas9 alone and dCas9-CtBP(L) led to about 20-25% repression of *Mpp6*. Targeting dCas9-CtBP(S) led to significant repression (∼50%), and this repression is greater than effects by dCas9-CtBP(L) alone, indicating a short isoform-specific effect on this gene. Values within the bars reflect the average relative expression. Controls are wing discs expressing non-targeting QUAS gRNA with *nubbin*-*GAL4*-driven dCas9, used to normalize all other samples. * p<0.05, ** p<.01, *** p<.001. Error bars indicate SD. Where the result of statistical analysis is not indicated, the value of p>0.05 (values indicated in **Supplementary Table 4**).

In this system, we did not find complete suppression of gene expression as noted in other transcriptional assays. However, an important caveat is that the level of repression measured may be an underestimate, because the *nubbin* driver is expressed only in the wing pouch, while we used the entire wing disc for RT-qPCR analysis. Interestingly, although CtBP(L) repressed *E2F2* and *Mpp6* to the same extent as dCas9 alone, it clearly showed more pronounced phenotypic effects in the adult stage. It may be that the CtBP(L) does have some specific activity in this setting (the trend, though not statistically significant, was slightly stronger than dCas9 alone). Alternatively, CtBP(L) may exert an effect later in development that we don’t measure at this timepoint. It is likely that the overt structural defects noted in the adult wing reflect perturbations to the complex gene regulatory networks that control wing development, which a single transcriptional measurement cannot capture.

### Position-sensitive CtBP repression in cell culture

Many tests of CtBP function have relied on transiently transfected reporter genes; however, few studies have directly compared repression activity on the same genes in their endogenous chromosomal location. To further assess CtBP(L) and CtBP(S) function and compare our *in vivo* results to traditional reporter assays, we expressed the dCas9-CtBP chimeras in S2 cells, using an *Mpp6* reporter which we have previously demonstrated is susceptible to repression by dCas9-Rb proteins [19]. Here, we employed seven individual gRNAs to test for possible position effects on this 1 kb promoter region (**Figure 4A**). Both CtBP(S) and CtBP(L) showed strongest effects with gRNA 2 and 5; dCas9 alone did not mediate significant repression from the gRNA 2 position, but did from gRNA 5, likely due to steric effects (**Figure 4B-D**). The dCas9 control did not mediate repression from any other site, clearly different from the CtBP effects with gRNAs 1, 2, and 3. A simple distance effect, with stronger repression proximal to the transcriptional start site, was not evident. Additionally, CtBP(S) appeared to be more effective at the more distal gRNA 1 and B positions than near the TSS, at 4. Overall, it is striking that CtBP(L) performed similarly to CtBP(S) on this reporter, given the clear differences noted for activity in the native chromosomal context.

**Figure 4.**
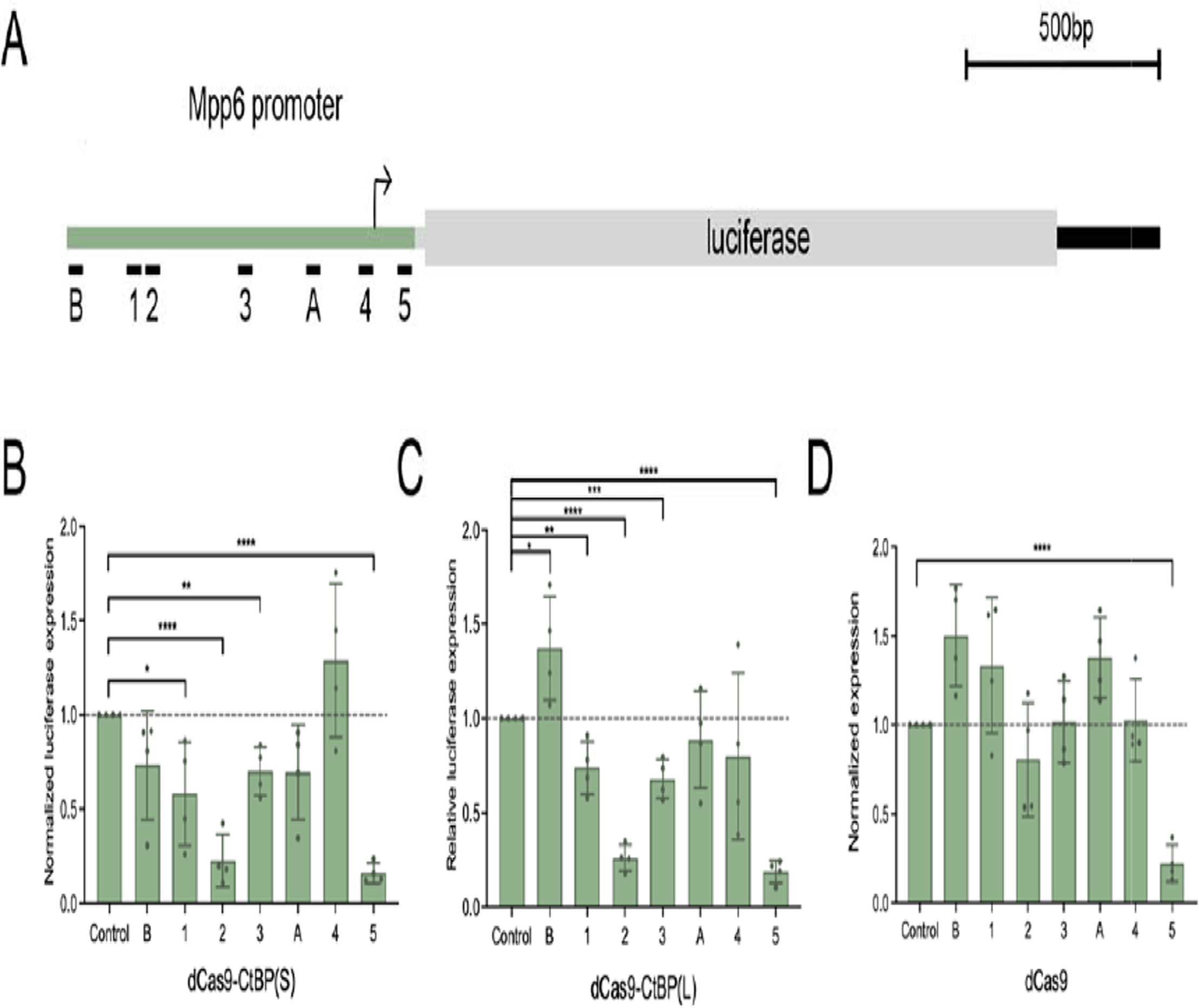
Testing Cas9-CtBP range of action on a luciferase reporter gene in S2 cells. S2 ce ls were transfected with *actin*-*GAL4*, the *Mpp6*-luciferase reporter, one of the dCas9 effectors, and a single gRNA. **A)** Schematic of luciferase reporter that was designed to be regulated by the *Mpp6* promoter, with gRNA positions indicated below. **B)** dCas9-CtBP(S) has position-specific effects. Position 2 led to the most severe repression. Position 5 caused the same level of repression as dCas9 alone, suggesting steric hindrance. **C)** dCas9-CtBP(L) has position-specific effects, which are similar to those of CtBP(S). **D)** The dCas9 control did not lead to significant repression, aside from position 5. The dCas9 results are the same control experiments as presented in [19]. Positions 4 and 5 are targeting sites for gRNAs at -577 and -672 shown in previous figures. Sequences of all gRNAs are described in [19]. * p<0.05, ** p<.01, *** p<.001, **** p<.0001. Error bars indicate SD.

## DISCUSSION

Here, we created novel dCas9-chimeras to CtBP corepressor proteins to compare their impact on gene expression in an *in vivo* system. Our study of Drosophila CtBP(L) and CtBP(S) isoforms using this CRISPRi approach has revealed that the two isoforms of this corepressor do exhibit different functional potential. Additionally, CtBP itself shows promoter selectivity, consistent with the findings of the Stark laboratory, where CtBP(S) was assayed against a wide spectrum of putative enhancers [16]. Our data suggest that CtBP proteins are involved in selective modulation of their gene targets, consistent with a “soft repression” form of regulation that may characterize many repressive interactions in the cell [23].

Evolutionary conservation of the CTD of CtBP indicates that this portion of the corepressor must be of importance; yet, most assays employed in previous studies have not identified a difference in function at the transcriptional level [6, 7]. One possible explanation is that the domain is involved in other aspects of CtBP biology, such as turnover or intracellular targeting, which may be overlooked in overexpression assays. Alternatively, its function in gene regulation may not have been identified yet, as the context in which CtBP has been assayed is limited; even the recent high throughput assessment of GAL4-CtBP(S) using STARR-seq was carried out with transient transfections and the significance of the CTD were not assessed [16]. There may be diverse roles for this IDR; however, our results strongly point to a transcriptional regulatory potential for the unstructured CtBP CTD.

Few studies have tested the impact of CtBP proteins with or without the conserved, long CTD on expression of endogenous genes, with the exception of genomic rescue experiments that demonstrated that viability is possible with either a CtBP(S) or CtBP(L) rescue construct [15]. However, the survivors from genomic rescues employing single isoforms showed a variety of phenotypes, including elevated embryonic lethality and aberrant wing development, indicating that limiting expression to one isoform alone does not fully satisfy developmental demands. Here, by directly testing CtBP isoforms using CRISPRi on endogenous genes in Drosophila, we uncovered a striking difference between CtBP(L) and CtBP(S). On the *Mpp6* promoter, CtBP(S) was a potent repressor of gene expression and caused a severe wing phenotype, while CtBP(L) was much milder in its transcriptional and phenotypic effects.

Many studies probing IDR functions have relied on high throughput assays to characterize IDRs en masse, and those focused on specific IDRs and proteins with disordered domains often use cell culture assays to uncover function. Therefore, it is important that with a direct comparison of our transcriptional effectors using transiently transfected reporters in S2 cells, we are unable to recapitulate the clear difference between CtBP(S) and CtBP(L) observed when targeting genes in a chromosomal context in the developing organism. Our finding that the CtBP(L) isoform is less active only on the chromatinized endogenous *E2F2/Mpp6* regulatory region provides support for the notion that the CTD regulation is chromatin-related. By combining an *in vivo* approach with CRISPRi, which is rarely done for dissecting mechanisms of gene regulation, we uncovered that the unstructured and highly conserved C-terminal domain of CtBP does in fact play a role in gene regulation. Additionally, our CRISPRi system ensures targeting to the promoter; thus, the CTD regulatory impact is likely to be at the level of transcriptional action, rather than promoter binding.

What might be the molecular action of this IDR on CtBP itself? Biochemical assays have shown that this intrinsically disordered domain is not required for NAD(H) binding or oligomerization—properties which are necessary for *in vivo* functionality [7, 23, 25]. The CTD of mammalian CtBP has been shown to be a target of post-translational modifications, including phosphorylation and sumoylation, which may affect conformation or protein-protein interactions of this domain [8]. It is interesting that a different eukaryotic dehydrogenase-like corepressor, NPAC/GLYR1, similar to CtBP, forms tetramers and possesses an IDR that is involved in functional contacts with histone-modifying lysine demethylases [26]. A similar function for the CtBP CTD may be uncovered in the future, but deeper understanding will require further biochemical and molecular genetic studies.

## EXPERIMENTAL PROCEDURES

### Plasmids used in this study

To create UAS:dCas9-CtBP constructs, the FLAG-tagged (DYKDDDDK) coding sequences for CtBP(L) and CtBP(S) were used, as described previously [14]. These coding sequences were amplified from their parent vector using 5’ *Pac*I and 3’ *Xba*I sites, and inserted in place of Rbf1 in the UAS:dCas9-Rbf1 plasmid described previously [19]. CtBP(L) is isoform F and CtBP(S) is a combination of isoform E and J, based on Flybase nomenclature. The *Mpp6*-luciferase reporter construct uses the *Mpp6* promoter, which includes the *Mpp6* 5’UTR, to drive luciferase expression, as was described previously [19]. The gRNA plasmids used in transfections were described previously, and target different sites of the *E2F2/Mpp6* bidirectional promoter [19].

### Transgenic flies

Flies were fed on standard lab food (molasses, yeast, corn meal) and kept at room temperature in the lab, under standard dark-light conditions. The *nubbin*-*GAL4* fly line was obtained from the Bloomington Drosophila Stock Center (BDSC; #25754) and was maintained as a homozygous line with a Chr 3 balancer obtained from BDSC #3704 *(w[1118]/Dp(1; Y)y[+]; CyO/Bl[1]; TM2, e/TM6B, e, Tb[1]*). Homozygous UAS:dCas9-CtBP flies were generated by using the □C31 integrase service at Rainbow Transgenic Flies Inc. #24749 embryos were injected with each dCas9-CtBP construct to integrate into Chr 3, landing site 86Fb. Successful transgenic flies were selected through the mini-*white* selectable marker expression in-house, and maintained as a homozygous line with Chr 2 balancer (from BDSC #3704). *nubbin*-*GAL4* and UAS:dCas9-CtBP homozygous flies were crossed to generate double homozygotes (*nubbin*-*GAL4*>UAS:dCas9-CtBP), using the Chr 2 and Chr 3 balancers (from #3704) These flies are donated to the Bloomington Drosophila Stock Center, and fly line numbers are indicated in **Supplementary Table 3**. sgRNA fly lines were obtained from the BDSC (fly line numbers indicated in **Supplementary Table 1**). Homozygous *nubbin-GAL4*>UAS:dCas9-CtBP flies were crossed to homozygous gRNA flies to generate triple heterozygotes (-/-; *nubbin*-*GAL4*/sgRNA; UAS:dCas9-CtBP/+) that are used for all fly experiments described here.

### Genotyping flies

All flies generated in this study were genotyped at the adult stage. Flies of each genotype were homogenized (1 fly/tube) in 50 μl squish buffer (1M Tris pH 8.0, 0.5M EDTA, 5M NaCl with 1μl of 10 mg/mL Proteinase K for each fly). Tubes were incubated at 37□ for 30 minutes, 95□ for 2 mins, centrifuged at 14,000 RPM in an Eppendorf Centrifuge 5430 for 7 minutes, and stored at 4□. Following PCR amplification, amplicons were cleaned using Wizard SV-Gel and PCR Clean-Up System, and sent for Sanger sequencing.

### Imaging adult wings

Adult wings were collected from ∼50 male and female 1-3 day-old adults. They were stored in 200 proof ethanol in -20□ until mounted. Wings were removed, mounted onto Asi non-charged microscope slides using Permount, and photographed with a Canon PowerShot A95 camera mounted onto a Leica DMLB microscope. Images were all taken at 10X magnification and using the same software settings.

### Wing disc dissections and RT-qPCR

50 third instar wing discs per biological replicate were dissected from L3 larvae and placed in 200 μl Trizol (Ambion TRIzol Reagent) and stored in -80□ until use. RNA was extracted using chloroform and the QIAGEN maXtract High Density kit, and stored in -80□. cDNA synthesis was performed using applied biosystems High Capacity cDNA Reverse Transcription Kit. RT-qPCR was performed using SYBR green (PerfeCTa SYBR Green FastMix Low ROX by Quantabio) and measured using the QuantStudio 3 machine by applied biosystems. Two control genes were measured and averaged (*Rp49, RpS13*) for all samples to provide a normalization standard to account for differences in RNA content in individual samples. To assess specific effects of targeted dCas9 proteins on promoters, *E2F2* and *Mpp6* levels were also measured in control wing discs obtained from crossing dCas9 to a non-targeting gRNA (QUAS). Primers used are found in **Supplementary Table 2**. RT-qPCR was performed on 3 biological replicates with two technical duplicates. Student’s t-test (two tailed, p<0.05) was used to measure statistical significance. Error bars indicate SD.

### Luciferase reporter assays

Reporter assays were performed as described previously, but with dCas9-CtBP(L) and dCas9-CtBP(S) effectors used here [19].

### Western blot

Drosophila S2 cells were grown in 25□ in Schneider Drosophila medium with glutamine (Gibco) containing 10% FBS and 1% penicillin-streptomycin (Gibco). 1.5 million cells were co-transfected with Effectene Transfection Reagent (Qiagen), according to manufacturer’s protocol. 250 ng of *actin*-*GAL4* (Addgene #24344) and 250 ng of UAS:dCas9-CtBP effectors were co-transfected in 6-well plates. Cells were harvested 3 days post-transfection and lysed using S2 lysis buffer (50mM Tris, pH 8.0; 150 mM NaCl; 1% Triton X-100), followed by boiling with Laemmli buffer. 100 μg of cell lysates were separated on a 4-20% resolving gel (Bio-Rad Mini-PROTEAN TGX Precast Gel #456-1094), transferred to a PVDF membrane for analysis using anti-FLAG (Sigma Aldrich #F3165, 1:10,000), and anti-CtBP (DNA208; [13]). Blocking with both primary and secondary antibodies was performed in 5% milk-TBST (500mM Tris-HCl, pH 7.4, 150 mM NaCl, 0.1% Tween 20). Blots were developed using HRP-conjugated GaM and GaR secondary antibodies (Pierce), and imaged using SuperSignal West Pico chemiluminescent substrate.

## ACKNOWLEDGEMENTS

We would like to thank members of the Arnosti lab for their thoughtful suggestions, and the Michigan State University RTSF and plasmidsaurus for sequencing plasmids.

## FUNDING

This work was supported by the National Institute for General Medical Sciences grant (number *R01GM124137)* to D.N.A., the National Institute of Child Health and Development grant (number *F31HD105410)* to A.M.R., and the BEACON luminaries grant to M.S.

## SUPPORTING INFORMATION

This article contains supporting information.

## CONFLICT OF INTEREST

The authors declare that they have no conflicts of interest with the contents of this article.

## DATA AVAILABILITY

The data supporting the findings of this study are available within the article and in the supplementary materials.

## Notes

### Competing Interest Statement

The authors have declared no competing interest.

### Summary of Updates

This manuscript has been revised to describe the broad implications of studying the CtBP C-terminal IDR, with updated analysis and statistical tests performed on the data presented in Figure 3.

