## Supplementary materials for "A regulatory role for the unstructured C-terminal domain of the CtBP transcriptional corepressor"

**Supplementary Table 1.** Table indicating genes that were targeted by dCas9-CtBP(S), the positions of the gRNA binding sites, and the BDSC line number. Targeting most of these genes did not cause a severe wing phenotype such as what was seen for *E2F2/Mpp6*.

| **Gene** | **Function** | **gRNA distance from TSS** | **BDSC line number** |
| --- | --- | --- | --- |
| *E2F2* | Cell cycle regulator | -577, -672 | 78707 |
| *Mpp6* | M-phase phosphoprotein | -18, +57 | 78707 |
| *Acf* | Subunit of two ATP dependent nucleosome remodeling complexes. | -42, -228 | 80174 |
| *InR* | Insulin receptor | -112, -420 | 78685 |
| *Atx2* | Involved in eye development | -476, -213 | 77320 |
| *Rbf1* | Transcriptional repressor | -65, -442 | 80755 |
| *Atf3* | Activating transcriptional factor | -112, -283 | 80180 |
| *p53* | Transcription factor | -211, -327 | 80207 |
| *vang* | Establishes planar polarity in epithelia | -69, -332 | 79671 |
| *dad* | Inhibitory SMAD in dpp pathway | -294, -398 | 79923 |
| *Cad99c* | Cell-cell adhesion | -36, -135 | 78154 |
| *Arm* | Cell adhesion and wingless signaling | -376, -95 | 78647 |
| *Dtg* | gastrulation | -257, -133 | 76074 |
| *mip40* | Critical regulator of the cell cycle | -428, -315 | 80293 |
| *Ap* | Transcription factor | -378, -148 | 80215 |
| *Caz* | Locomotion and eye development | -455, -334 | 79806 |
| *Mnt* | Transcription repressor | -426, -339 | 79927 |
| *Cyc* | Transcription of circadian clock genes. | -249, -93 | 78706 |
| *Hh* | Signaling pathway ligand | -163, -114 | 67560 |
| *Wwox* | Oxidoreductase | -191, -359 | 77229 |
| *DNApola* | Catalytic subunit of DNAP | -57, -202 | 78146 |
| *Sta* | Ribosomal protein | -72, -302 | 78627 |
| *mRps22* | Mitochondrial ribosomal protein | -476, -364 | 78159 |
| *GstE13* | Glutathione | -471, -244 | 77283 |
| *spen* | Regulator of wnt signaling | -424, -355 | 80177 |
| *wg* | Encodes a ligand of the Wnt/Wg signaling pathway | -314, -366 | 67545 |
| *mcm6* | Subunit of the hetero-hexameric mcm complex | -438, -383 | 79845 |
| *dpp* | Ligand of the transforming growth factor β signaling pathway | -450, -3 | 67554 |
| *Rbf2* | Cell and development regulator | -410, -187 | 79982 |
| *elob* | Control of wing cell identity. | -374, -266 | 79926 |

**Supplementary Table 2. Primers used for RT-qPCR analysis.** F is for Forward primer and R is for Reverse.

| **Gene** | **Primers (Forward and Reverse)** |
| --- | --- |
| *Rp49* | F: ATCGGTTACGGATCGAACAAGC  R: GTAAACGCGGTTCTGCATGAGC |
| *Rps13* | F: GGTCGTATGCACGCTCCT  R: CATCTGCGTTCAGTTTCAGC |
| *E2F2* | F: GACGAGGAAGTAGATATCAAGCG  R: TCAAAGAACCCATCCACATCG |
| *Mpp6* | F: GCTCGGTCATTCTGCTTTTG  R: CTCGGCTTTGATTTGGATGG |

**Supplementary Table 3. Fly lines generated in this study and donated to BDSC.**

| **BDSC Fly line #** | **Genotype** | **As referenced in this study** |
| --- | --- | --- |
| 99963 | *w[1118]; M{RFP[3xP3.PB] w[+mC]=UAS-dCas9.FLAG}ZH-86Fb* | UAS:dCas9 |
| 99964 | *w[1118]; M{RFP[3xP3.PB] w[+mC]=UAS-dCas9-Ctbp.L}ZH-86Fb* | UAS:dCas9-CtBP(L) |
| 99965 | *w[1118]; M{RFP[3xP3.PB] w[+mC]=UAS-dCas9-Ctbp.S}ZH-86Fb* | UAS:dCas9-CtBP(S) |

**Supplementary Table 4. p-values from unpaired, two-tailed Student’s t-test of values in Figure 3.** Highlighted values were statistically significant p-values that are indicated in the graphs. Purple highlight corresponds to the two statistically significant p-values (<.05) indicated in Figure 3B, and green are those represented in Figure 3C.

| *E2F2 p-values* | compared to | | | |
| --- | --- | --- | --- | --- |
|  | **control** | **dCas9** | **CtBP(L)** | **CtBP(S)** |
| **control** |  | 0.0418 | 0.0204 | 0.285 |
| **dCas9** |  |  | 0.8697 | 0.1004 |
| **CtBP(L)** |  |  |  | 0.0891 |
| *Mpp6 p-values* | compared to | | | |
|  | **control** | **dCas9** | **CtBP(L)** | **CtBP(S)** |
| **control** |  | 0.0324 | 0.0059 | 0.0002 |
| **dCas9** |  |  | 0.3821 | 0.0057 |
| **CtBP(L)** |  |  |  | 0.0084 |

**SUPPLEMENTARY FIGURES**

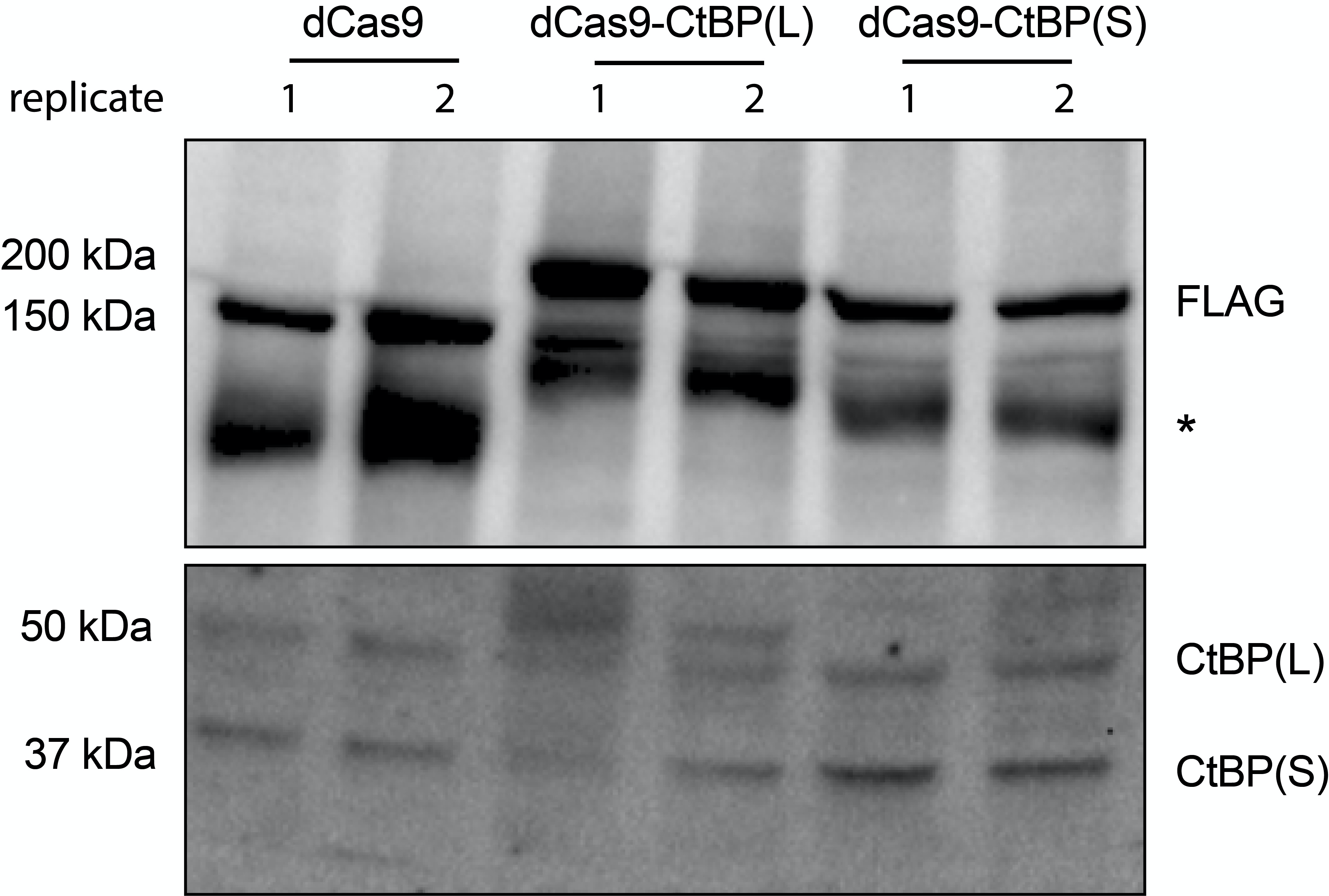

**Figure S1. dCas9-CtBP effectors are expressed in S2 cells** **.** dCas9, dCas9-CtBP(L), and dCas9-CtBP(S) were co-transfected in S2 cells with *actin*-*GAL4* in duplicate. Effectors run at expected sizes of ~150 kDa for dCas9 alone, and <200 kDa for the CtBP effectors. dCas9-CtBP(S) runs faster than dCas9-CtBP(L), consistent with a size difference of ~10 kDa. Asterisk (*) indicates presumptive degradation product. CtBP(L) and CtBP(S) (bottom panel) are the loading control for endogenous CtBP for these samples, using an anti-CtBP antibody (DNA208; [13]).

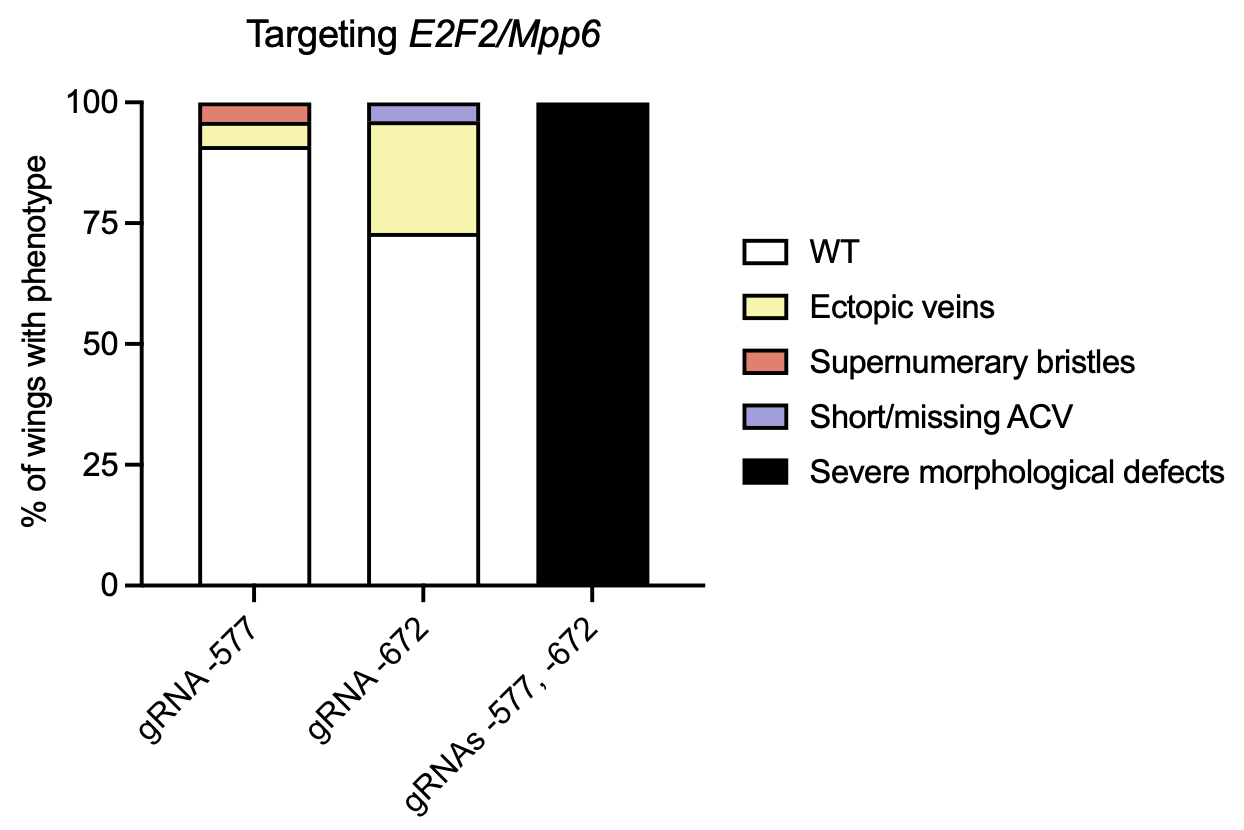

**Figure S2. Single gRNAs produce milder CtBP(S) effects.** Recruiting dCas9-CtBP(S) with individual gRNAs at -577 or -672 led to much milder effects than when used together in tandem, suggesting a possible cooperative effect when two dCas9-CtBP(S) molecules are brought together.
